## Supplementary Material for "Influences of time of day on generalization"

| | Number of Sessions | Delay Between Sessions | Session 1 Start Time | Session 2 Start Time | Bridge Exposure | Exemplars Per Category | Number of Test Trials | Order of Tasks |
|---|---|---|---|---|---|---|---|---|
| **Exp. 1** | 2 | 12 hours | Morning: 9AM Evening: 8PM | Morning: 9:30PM Evening: 9AM | To criterion | 5 | 53 | |
| **Exp. 2** | 2 | 12 hours | Morning: 9AM Evening: 8PM | Morning: 9:30PM Evening: 9AM | Morning: 1 block Evening: 6 blocks | 5 | 53 | |
| **Exp. 3** | 2 | 24 hours | Morning: 9AM Evening: 8PM | Morning: 9AM Evening: 8PM | To criterion | 5 | 53 | |
| **Exp. 4** | 1 | | 4PM | | To criterion | 5 | 53 | |
| **Exp. 5** | 1 | | Morning: 9AM Evening: 8PM | | To criterion | 3 | 21 | RAT, Distractor Priming, Generalization |
| **Exp. 6** | 1 | | Morning: 9AM Evening: 8PM | | To criterion | 3 | 21 | Generalization, RAT, Distractor Priming |
| **Exp. 7** | 1 | | Morning: 9AM Evening: 8PM | | To criterion | 5 | 53 | Generalization, RAT, Distractor Priming |

**Supplementary Table 1.** Experiment design differences. Refer to the Methods section for detailed protocol for each experiment. **Number of Sessions** = number of experimental sessions; two session experiments reflect consolidation-related designs wherein a post-delay test was administered in Session 2. **Delay Between Sessions** = approximate length of the delay in hours between Session 1 and Session 2. **Session 1 Start Time** = approximate start time of Session 1 for the Morning and Evening group. **Session 2 Start Time** = approximate start time of Session 2 (referred to in text as Test 2) for the Morning and Evening group. **Bridge Exposure** = total number of blocks (10 trials per block) participants encountered bridge satellites during the bridge training phase. Criterion was reached when a participant achieved nine out of ten trials in a block correct. Note that in Experiment 2 the number of blocks are fixed differently for each group, reflecting our aim to match initial generalization performance in this experiment. **Exemplars Per Category** = the number of unique satellites in each of the three categories encountered during the training phase. Number of Test Trials = total number of trials during the test phase in one session. **Order of Tasks** = the order of the tasks that participants completed in experiments that administered additional measures (RAT, Distractor Priming). Generalization refers to the full generalization paradigm (i.e. both training and test).

| | | Number of Blocks to Criterion | | Feature Accuracy Last Block | | Class Accuracy Last Block |
|---|---|---|---|---|---|---|
| | | **Initial** | **Bridge** | **Initial** | **Bridge** | **Initial** |
| **Exp. 1** | Morning | 4.15±0.36 | 2.46±0.27 | 0.73±0.02 | 0.95 ±0.01 | 0.94±0.03 |
| | Evening | 4.94±0.50 | 2.59±0.32 | 0.73±0.02 | 0.94±0.01 | 0.96±0.01 |
| | Group Diff. *p* | .24 | .77 | .96 | .56 | .63 |
| **Exp. 2** | Morning | 4.36±0.48 | 1 (fixed) | 0.71±0.02 | 0.69±0.06 | 0.98±0.01 |
| | Evening | 3.77±0.46 | 6 (fixed) | 0.73±0.03 | 0.98 ±0.01 | 0.98±0.01 |
| | Group Diff. *p* | .38 | | .59 | .000005*** | .61 |
| **Exp. 3** | Morning | 4.07±0.32 | 2.71±0.27 | 0.73±0.02 | 0.94±0.01 | 0.96±0.01 |
| | Evening | 4.38±0.27 | 2.54±0.31 | 0.76±0.01 | 0.96±0.01 | 0.97±0.01 |
| | Group Diff. *p* | .46 | .67 | .38 | .35 | .56 |
| **Exp. 4** | Nap | 4.45±0.44 | 2.10±0.14 | 0.76±0.02 | 0.97±0.01 | 0.98±0.01 |
| | No Nap | 3.94±0.39 | 2.52±0.23 | 0.68±0.04 | 0.95±0.01 | 0.96±0.02 |
| | Group Diff. *p* | .40 | .11 | .07 | .30 | .65 |
| **Exp. 5** | Morning | 4.36±0.41 | 2.28±0.11 | 0.77±0.02 | 0.95±0.01 | 0.98±0.01 |
| | Evening | 4.39±0.45 | 1.94±0.17 | 0.76±0.02 | 0.97±0.01 | 0.98±0.01 |
| | Group Diff. *p* | .96 | .09 | .72 | .35 | .75 |
| **Exp. 6** | Morning | 4.12±0.33 | 2.31±0.16 | 0.74±0.01 | 0.96 ± 0.01 | 0.97±0.01 |
| | Evening | 5.57±0.47 | 2.52±0.23 | 0.76±0.02 | 0.95 ± 0.01 | 0.96±0.02 |
| | Group Diff. *p* | .01* | .43 | .17 | .50 | .77 |
| **Exp. 7** | Morning | 4.34±0.22 | 2.17±0.11 | 0.68±0.02 | 0.95±0.01 | 0.96±0.01 |
| | Evening | 3.74±0.21 | 2.21±0.11 | 0.70±0.02 | 0.97±0.01 | 0.94±0.02 |
| | Group Diff. *p* | .05* | .80 | .67 | .07 | .31 |

**Supplementary Table 2.** Training performance. Measures of performance during the main training phase (Initial) and the following bridge training phase (Bridge) are shown for each experimental group. Descriptive statistics (mean ± one SEM) and group differences (*p*-values from t-test) are shown. Note that the number of trials per block in the initial training phase was lower in Exp. 5 and Exp. 6. ***$p < .001$, *$p < .05$.

| | | MEQ Score | MEQ Chronotype Proportions (Morning /Neutral/ Evening) | SSS | H Since Wake | Sleep Qual. Last Night | H Asleep Last Night | Epworth |
|---|---|---|---|---|---|---|---|---|
| **Exp. 1** | Morning | | | 2.85±0.22 | 2.01±0.21 | 2.00±0.20 | 6.92±0.19 | 6.92±1.12 |
| | Evening | | | 2.35±0.15 | 11.77±0.40 | 2.06±0.16 | 8.35±0.34 | 5.53±0.72 |
| | Group Diff. *p* | | | .06 | | .82 | .002** | .29 |
| **Exp. 2** | Morning | | | 2.36±0.23 | 1.76±0.16 | 1.64±0.20 | 7.66±0.32 | 6.07±0.66 |
| | Evening | | | 2.61±0.27 | 12.22±0.32 | 2.15±0.25 | 7.56±0.38 | 7.85±1.07 |
| | Group Diff. *p* | | | .46 | | .12 | .83 | .16 |
| **Exp. 3** | Morning | 47.93±2.65 | 21% / 50% /29% | 2.14±0.18 | 1.93±0.16 | 2.07±0.16 | 7.35±0.32 | 5.50±0.87 |
| | Evening | 50.46±2.65 | 15% / 70% /15% | 3.23±0.43 | 13.01±0.86 | 1.92±0.14 | 7.33±0.36 | 5.69±0.93 |
| | Group Diff. *p* | .51 | | .02* | | .50 | .96 | .88 |
| **Exp. 4** | Nap | 46.90±2.46 | 10% / 60% / 30% | 2.50±0.18 | 8.02±0.26 | 1.85±0.13 | 7.40±0.30 | 7.20±0.63 |
| | No Nap | 48.65±2.47 | 24% / 47% / 29% | 2.35±0.17 | 7.81±0.43 | 2.12±0.17 | 7.60±0.27 | 6.47±0.51 |
| | Group Diff. *p* | .62 | | .58 | .66 | .21 | .62 | .39 |
| **Exp. 5** | Morning | 49.08±1.55 | 8% / 72% /20% | 2.00±0.12 | 1.94±0.16 | 1.80±0.16 | 7.09±0.15 | 5.92±0.55 |
| | Evening | 51.56±2.11 | 17%/ 67% /17% | 2.44±0.25 | 13.10±0.23 | 2.06±0.19 | 7.77±0.25 | 6.06±0.86 |
| | Group Diff. *p* | .34 | | .08 | | .31 | .02* | .89 |
| **Exp. 6** | Morning | 49.81±1.68 | 15% / 69% / 15% | 2.08±0.17 | 2.02±0.18 | 1.88±0.14 | 7.51±0.20 | 6.62±0.65 |
| | Evening | 50.55±2.30 | 14% / 59% / 27% | 2.68±0.20 | 13.43±0.27 | 1.73±0.15 | 7.82±0.15 | 4.95±0.62 |
| | Group Diff. *p* | .79 | | .03* | | .45 | .23 | .07 |
| **Exp. 7** | Morning | 48.29±1.23 | 10% / 63% / 27% | 2.22±0.12 | 1.77±0.09 | 1.73±0.09 | 7.41±0.12 | 6.46±0.60 |
| | Evening | 45.81±1.22 | 3% / 76% / 22% | 2.38±0.17 | 12.25±0.21 | 1.76±0.10 | 7.86±0.15 | 6.32±0.55 |
| | Group Diff. *p* | .16 | | .45 | | .85 | .03* | .86 |

**Supplementary Table 3.** Survey measures. Group descriptive statistics (mean ± one SEM) and Morning and Evening group comparisons (t-tests) for each experiment are shown. For two session experiments (Experiment 1, Experiment 2, Experiment 3) only Session 1 survey responses are presented. **MEQ Score =** score on the Morningness-Eveningness Questionnaire, with higher values reflecting a circadian preference towards the morning, **MEQ Chronotype Proportions =** assigned chronotype using the standard Morningness-Eveningness Questionnaire scoring, percentages are rounded to the nearest whole number, **SSS** = participant response on the Stanford Sleepiness Scale, with higher values indicating more sleepiness, **H Since Wake** = number of hours since wake time in the morning (calculated from participant reported wake time), **Sleep Qual. Last Night** = participant reported quality of sleep the night prior to the experiment (1 = Excellent, 2 = Good, 3 = Fair, 4 = Poor.), **H Asleep Last Night =** participant reported hours spent asleep the night prior to the experiment, **Epworth =** score on the Epworth Sleepiness Scale, with higher values indicating more daytime sleepiness.**$p < .01$, *$p < .05$. In Experiment 3, the Evening group in this sample was sleepier than the Morning group. This difference did not relate to generalization in regression models (*p*'s > .16).

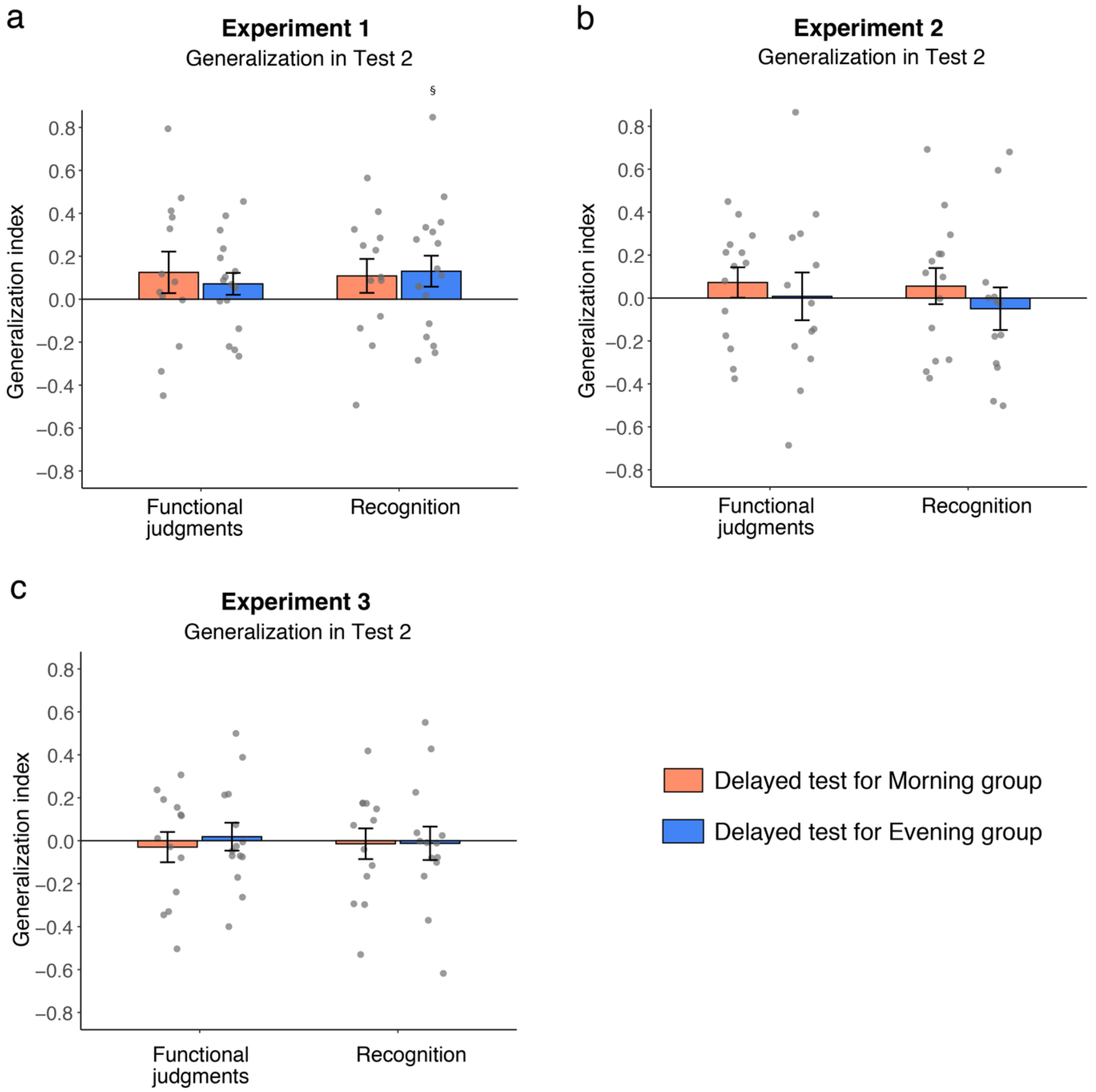


**Supplementary Figure 1.** Generalization in Test 2 for **(a)** Experiment 1, **(b)** Experiment 2, and **(c)** Experiment 3. For Experiments 1 & 2 (12h delay), the Morning group was tested in the morning in Test 1 and evening in Test 2, and the Evening group was tested in the evening in Test 1 and morning in Test 2. For Experiment 3 (24 hour delay), the Morning group was tested in the morning on both Test 1 and Test 2, and the Evening group was tested in the evening on both Test 1 and Test 2.

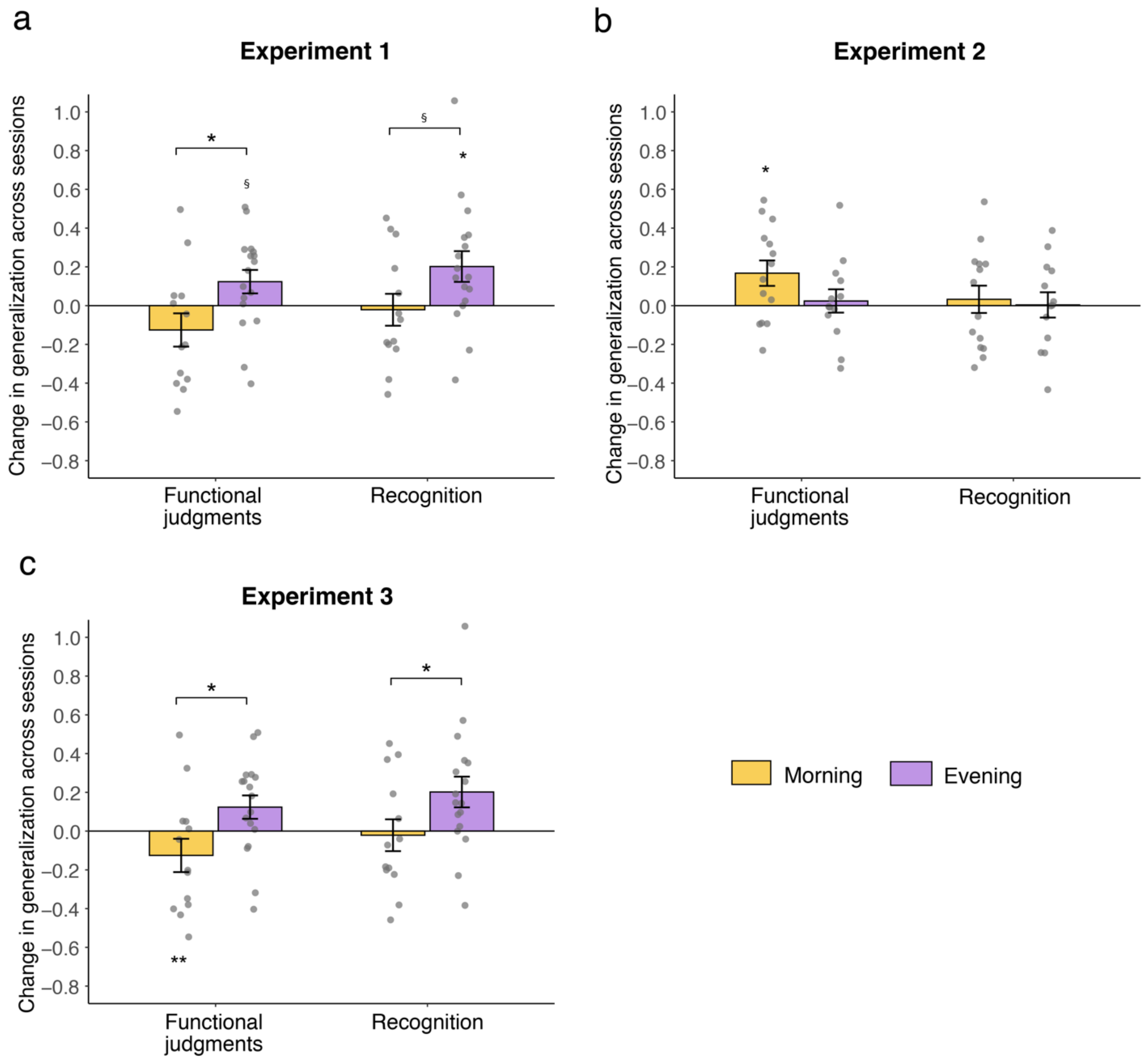


**Supplementary Figure 2.** Change in generalization across sessions (Test 2 Generalization – Test 1 Generalization) for **(a)** Experiment 1, **(b)** Experiment 2, and **(c)** Experiment 3. For Experiments 1 & 2 (12h delay), the Morning group was tested in the morning in Test 1 and evening in Test 2, and the Evening group was tested in the evening in Test 1 and morning in Test 2. For Experiment 3 (24 hour delay), the Morning group was tested in the morning on both Test 1 and Test 2, and the Evening group was tested in the evening on both Test 1 and Test 2. *$p < .05$, §$p < .1$

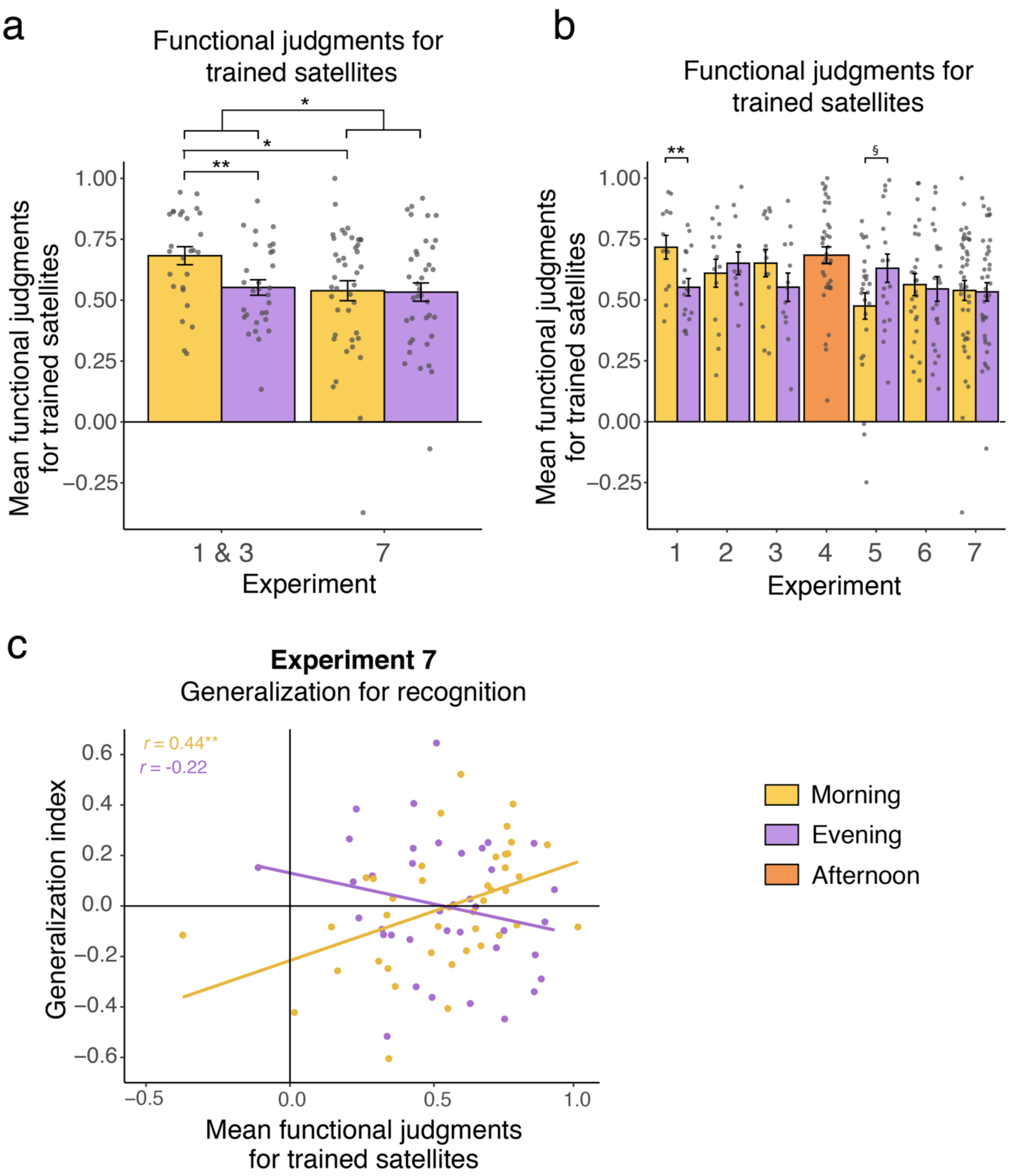


**Supplementary Figure 3.** Functional judgments for trained satellites. **(a)** Functional slider judgments for trained satellites in Experiment 1&3 (pooled) and Experiment 7. **(b)** Functional slider judgments for trained satellites across all experiments. The reliable Morning vs. Evening group differences within experiments are marked. **(c)** Interaction between time of day and functional judgments for trained satellites and generalization for recognition in Experiment 7. Post-hoc Pearson's correlations were computed separately for the Morning and Evening group and are depicted in the top left quadrant. *$p < .05$, ** < .01 §$p <$ .1

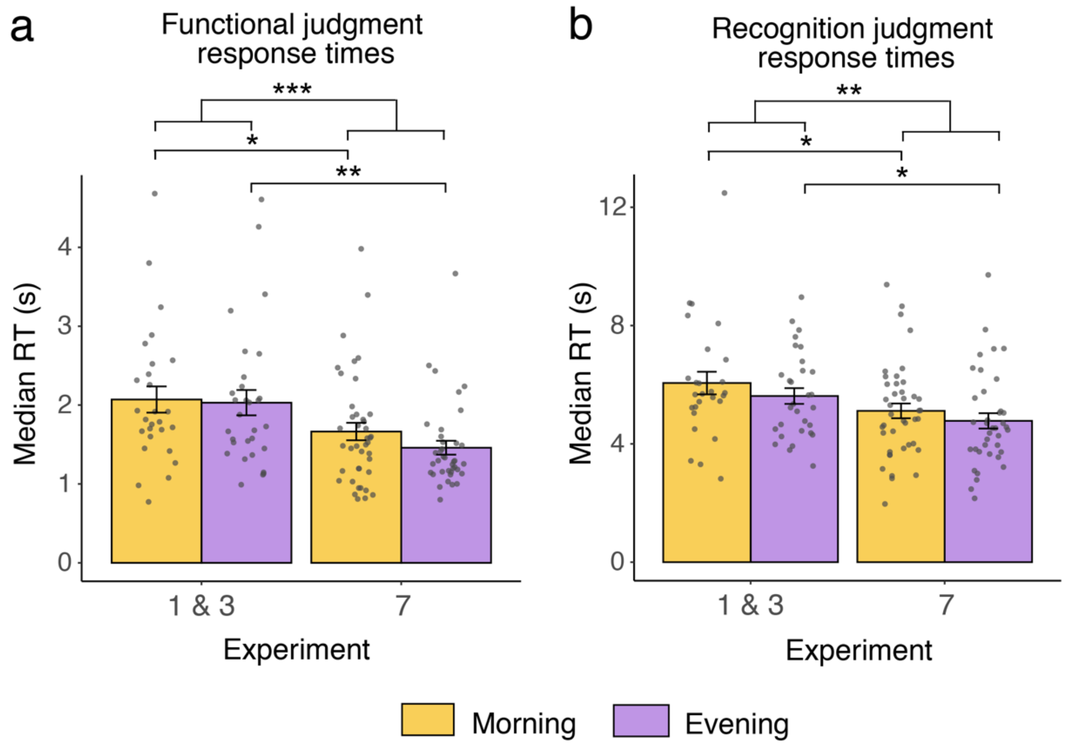


**Supplementary Figure 4.** Overall slider response times. Median slider response time differences in Experiments 1 and 3 (pooled; E1&3) versus Experiment 7 (E7) for **(a)** functional judgments and **(b)** recognition judgments. ***$p < .001$, **$p < .01$, *$p < .05$.
