## Supplementary material for "Influences of time of day on generalization": read_me.txt

test_data.csv 

Contains the test data for Experiments 1-7. Participant RAT and Distractor Priming scores, as well as survey measures are also appended.

subject: subject ID
experiment:  experiment number (1-7)
group: experimental group (morning or evening, nap or no nap)
session: first or second test session (1,2)

trial: trial number
trained: whether the satellite was shown during any part of training
bridge_type: whether the bridge-satellite was consistent with the categories connected during the bridge training phase
feature_type: the category typicality of the features that made up the bridge satellite
trained_type: whether or not the trained satellite was seen during initial training or bridge training

seen_slider_resp: slider response for the recognition judgment
seen_slider_rt: slider response time (seconds) for the recognition judgment
func_slider_resp: slider response for the functional judgment
func_slider_rt: slider response time (seconds) for the functional judgment
code_name_corr: whether the code name was correct
code_name_rt: code name response time (seconds)

rat: remote associates test score for that participant
priming: distractor priming score for that participant

meq: morningness-eveningness questionnaire score (a higher value indicates greater morningness)
sss: stanford sleepiness score in session 1 (1 = “Feeling active, vital, alert, or wide awake”, 8 = “Asleep”)
sss_sess2: stanford sleepiness score in session 2
h_asleep: self-reported hours asleep the night prior to session 1
sleep_qual: self-reported sleep quality the night prior to session 1
epworth: epworth sleepiness scale (a higher value indicates greater daytime sleepiness)

For training data, excluded participants, or other information, please
